# Variants in Cas9 and *nanos* regulatory elements modulate activity and reduce resistance allele formation in homing gene drive

**DOI:** 10.1101/2025.11.04.686506

**Authors:** Ruizhi Zhou, Jie Du, Nicky R. Faber, Jackson Champer

## Abstract

Gene drive is a novel approach for controlling vector borne disease via either population modification or suppression. Even with high efficiency, though, overall drive performance can be reduced by somatic Cas9 expression and by maternal deposition of Cas9, leading to resistance allele formation. The *nanos* promoter for Cas9 shows very little leaky somatic expression, but it causes high rates of embryo resistance allele formation in *Drosophila melanogaster*. By truncating the promoter, we reduced rates of embryo resistance to undetectable levels, but germline cutting in females decreased by over half. Germline cutting and successful drive conversion was eventually lost when only the 5′ UTR was present, though males still retained moderate germline drive efficiency. Several additional methods were tested to improve performance, including additional suppressor elements to the 3′ UTR and introns to increase expression level. The most successful of these was the addition of a second nuclear localization signal, which substantially increased activity when coupled with a full-length or truncated *nanos* promoter. Overall, these experiments show the potential to modulate Cas9 regulatory elements to achieve desired expression for gene drive applications, while also showcasing the difficulty of obtaining an optimal activity profile.

## Introduction

Gene drives employ super-Mendelian inheritance to achieve genome editing at a population scale^1–5^. They can generally be divided into modification and suppression drive types. Modification drives can help to spread a desirable trait across a population, including reducing pesticide resistance or reducing infectious disease transmission from a vector species. Suppression drives target an essential gene and can eventually collapse a population. Within these categories, there are many subtypes of gene drives with different levels of confinement. Although CRISPR toxin-antidote gene drives^6–11^ can be confined to a target population and are easier to engineer, homing gene drives that copy themselves are substantially more powerful and may still be an important part of a confined suppression strategy^12–14^. It is thus critical to develop methods for engineering efficiently performing homing gene drive.

In CRISPR homing gene drives, the Cas9 complex first binds to the gRNA target site and then induces a double-strand break^1–5^. Homology-directed repair subsequently uses the drive allele as a repair template, copying it into the wild-type allele. However, end-joining is another major DNA repair pathway that directly joins two ends of the break together, often mutating the sequence and preventing future recognition by the gRNA. These mutations are termed resistance alleles and can greatly affect the success of the gene drive. Functional resistance alleles can outcompete the gene drive, which happens quickly for suppression drives as these have recessive fitness effects by design. Even if steps are taken to prevent functional resistance alleles from forming, suppression drives can also be substantially affected by nonfunctional resistance.

*Drosophila melanogaster* is a model organism that has been widely studied in the field of gene drive because it has many genes that have homologs in multiple agricultural pests and disease vectors. Currently, homing gene drives are the most advanced in *Anopheles* malaria mosquitoes, usually reaching inheritance rates of 95-100%^15–19^. This potentially already allows the construction of powerful suppression and effective modification gene drives. However, in many species, including *Drosophila melanogaster*^20–22^, *Drosophila suzukii*^23,24^, *Ceratitis capitata*^25^, *Mus musculus*^26^, *Rattus* sp.^27^, *Plutella xylostella*^28^, and *Aedes aegypti*^29,30^, homing gene drives still face considerable limitations due to varying combinations of limited drive conversion rate, high resistance allele formation, high somatic expression, or low Cas9 cleavage rates. A low conversion rate can slow population modification or even prevent it if the drive has substantial fitness costs. Both low drive conversion and high somatic expression are particularly detrimental to suppression drives^31^.

With gRNA expression having limited flexibility, the main way to control Cas9 cleavage patterns is by modulating Cas9 regulatory elements (promoter, 5′ UTR, 3′ UTR) or the nuclease itself. An ideal promoter has high expression in the germline to promote drive conversion, while having low maternal deposition into embryos (as this causes resistance allele formation) and minimal somatic expression. Recently, several different promoters were examined in fruit flies^32^. *CG4415* was successful in suppressing a population, but it still induced some somatic expression and likely has lower overall activity. The *nanos* promoter, on the other hand, had no detectable somatic expression and high germline expression, though maternal deposition was also high. Furthermore, *nanos*-based gene drives were even more effective in *Drosophila suzukii*^23^ and *Anopheles gambiae*^33,34^, making it a promising candidate for further development. Notably, the *D. suzukii nanos*-Cas9 element showed superior performance in *D. suzukii* compared to the same system only with sequences based off the *D. melanogaster nanos* gene, but this may also have been because the *D. suzukii* sequence Cas9 had a second nuclear localization signal (NLS), which may increase cleavage activity^23^. Several other studies provided clues that may be useful for adjusting Cas9 activity. Some reports indicated that a Syn21 intron^35^ or improved Kozak sequence^36^ could also increase expression. A report on a truncated *nanos* promoters indicated that germline activity can be preserved while losing maternal deposition^37,38^. At the 3′ UTR, previous studies showed that a particular loop element may be important for suppressing the translation of maternally deposited mRNA in the early embryo, providing a possible strategy to reduce the effect of maternal deposition^37,38^.

The objective of this study is to further explore how optimization of *nanos* regulatory elements and Cas9 itself could adjust cleavage rates and eliminate maternal Cas9 deposition in the embryonic stage. We also examined the effect of a second NLS. We found that truncated versions of *nanos* remain effective, but a large enough truncation loses substantial activity, eliminating maternal deposition while still preserving moderate germline expression. Truncating the system to the 5′ UTR allows activity only in males. However, by including three copies of a short *nanos* promoter element in front of the 5′ UTR, full activity was restored. Similarly, the addition of a second NLS substantially improved activity. When both of these were combined, somatic expression was observed. Modifications involving addition of a Syn21 intron, consensus Kozak sequence, and modification of the 3′ UTR did not substantially alter promoter performance, while a PEST sequence eliminated most Cas9 activity. Overall, while we could not achieve optimal expression with *nanos*-Cas9, we were still able to adjust activity levels both upward and downward, allowing more control over gene drive performance for future applications.

## Methods

### Plasmid construction

For molecular cloning, restriction digest, Gibson Assembly, and PCR, reagents were obtained from New England Biolabs; 5-α competent *Escherichia coli* was obtained from TIANTEN and New England Biolabs; oligonucleotides were obtained from Integrated DNA technologies. ZymoPure Midiprep kit from Zymo Research were used to purify plasmids for egg microinjection. Plasmid sequences were confirmed by Sanger sequencing. We provide annotated sequences of the donor plasmids on GitHub in APE format^39^ (https://github.com/jchamper/Nanos-Cas9-Regulation).

### Fly rearing and generation of transgenic lines

Embryo injections were done by Fungene or Rainbow Transgenic Flies. *w*^*1118*^ flies were injected with donor plasmids (500 ng/µL), Cas9 helper plasmid TTChsp70c9^40^ (450 ng/µL) and gRNA helper plasmid BHDabg1^41^ (100 ng/µL). All flies were fed with modified Cornell standard cornmeal medium (with 10 g agar instead of 8 g per liter, addition of 5 g soy flour, and without the phosphoric acid) in vials within an incubator with a temperature of 25°C and on a 14/10-hour day/night cycle. Injected flies were crossed with *w*^*1118*^ flies, and their offspring were screened for fluorescence indicating successful knock-in. When screening transgenic lines, flies were anesthetized using a CO_2_ pad.

### Genotype and phenotype

Fluorescent genes *EGFP* and *DsRed* were expressed using the 3xP3 promoter, allowing for visualization of both fluorescent proteins in the eyes of the flies using NIGHTSEA adapters (SFA-GR for DsRed and SFA-RB-GO for EGFP). The presence of DsRed indicates that the fly strain contains a drive with gRNAs (that may have Cas9 if it targets EGFP), while EGFP shows that the fly strain contains a Cas9 gene or possesses only EGFP itself as a synthetic target site. The body phenotype is wild-type, except for lines where the gRNA targeted the X-linked gene *yellow*, where both the drive and nonfunctional resistance alleles recessively cause yellow body color. This can be used to visualize resistance alleles that form in the germline or early embryo. It can also be used to assess somatic drive activity through visible mosaicism. Some gRNA lines target the fertility genes *doublesex* (*dsx*) and *yellow-G*, which can give rise to recessive female-sterile phenotypes. One gRNA line targets the haplolethal gene *RpL35A*, and any resistance alleles are thus nonviable (the drive retains a rescue element, allowing any combination of drive and wild-type to remain viable).

### Crosses and data analysis

Cas9 lines were first crossed with gRNA lines to assess drive performance. Individuals with both green and red fluorescent eyes were then crossed with *w*^*1118*^, and they were allowed to lay eggs for seven days in individual vials (each containing one male and one female). Their offspring were then phenotyped and used to calculate drive performance. The drive inheritance rate was based on the percentage of offspring with DsRed eyes. Using a gene drive targeting the *yellow* gene, more information can be obtained; females with red and green eye fluorescence as well as yellow body phenotype after the first cross indicate somatic drive activity. Further, for female drive carriers, offspring without EGFP (indicating the absence of the Cas9 element) but with yellow body phenotype indicate resistance alleles that formed in the embryo from maternally deposited Cas9 and gRNA (the requirement for lacking EGFP/Cas9 can be dropped if somatic activity is minimal). Two gRNA elements targeting other genes were also similarly used to determine drive inheritance, but resistance alleles were not directly assessed. In another type of cross, a complete drive with DsRed is crossed to flies with a synthetic target EGFP, and the resulting progeny are also crossed to measure drive performance. Drive inheritance is the same as before. If female heterozygotes are crossed to males with EGFP, embryo resistance is indicated by progeny that lack the EGFP fluorescent phenotype. Germline resistance in male drive heterozygotes are present in offspring lacking EGFP if the males are crossed to females lacking EGFP.

### Data analysis

Each individual fly is considered as a separate batch because not every offspring from a single parent may have independently received a drive or resistance allele (Cas9 activity can occur in early germline cells that give rise to several offspring). To account for such batch effects, we analyzed our data as in previous studies^42^. We fit a generalized linear mixed-effects model with a binomial distribution (maximum likelihood, Adaptive Gauss-Hermite Quadrature, nAGQ = 25), enabling variance between batches by including a random effect per batch, which results in slightly different parameter estimates and higher standard error estimates than if each offspring were considered an independent sample from a binomial distribution. This analysis was conducted with R (3.6.1) using the packages lme4 (1.1-21) and emmeans (1.4.2).

## Results

### Cas9 gene elements

In this study, we sought to generate Cas9 elements with superior performance to achieve efficient homing gene drive. A previous study indicated that the *nanos* promoter was strong in the germline and had minimal somatic expression^32^. However, it also caused high maternal deposition and subsequent formation of embryo resistance alleles (Figure S1). Such resistance allele formation will have detrimental effects on gene drives, especially suppression systems, even if functional resistance is avoided by gRNA multiplexing and other methods. We chose our well-characterized *nanos*-Cas9 split element in *Drosophila melanogaster* as the basis for several variant designs to improve gene drive performance. Each Cas9 element contained an EGFP fluorescence gene expressed by the 3xP3 promoter (for expression in the eyes of adults) and terminated by an SV40 element (Figure 1). These were placed at two nearby genomic sites (downstream of two adjacent genes on chromosome 2R) with slightly different construct component orientations, but these differences likely did not affect performance^32^. Most testing involved a split gRNA element targeting the X-linked *yellow* gene^41^, allowing easy assessment of embryo resistance alleles, but only allowing the measurement of drive conversion in females (Figure S1). Some additional assessments were conducted with a split homing rescue drive targeting a haplolethal gene^22^ and a split homing suppression drive targeting a female fertility gene^43^ (Figure S1).

**Figure 1.**
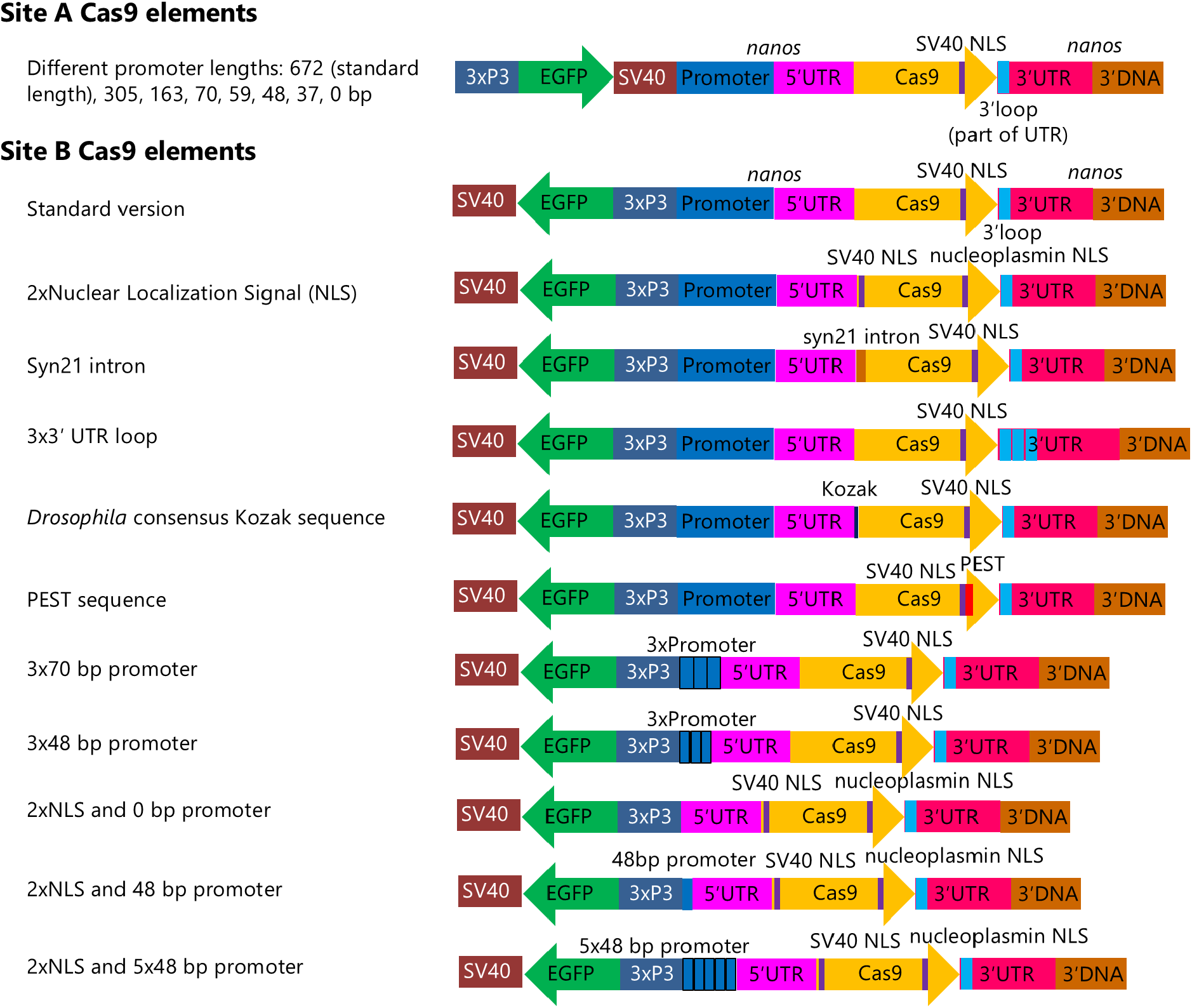
Cas9 element variants of this study. Each split Cas9 element contains an *EGFP* fluorescent marker gene driven by the 3xP3 promoter and terminated by a SV40 3′ UTR. At site A, the *EGFP* gene is in the same direction as Cas9, with eight different variants having different lengths of *nanos* promoter (the distal end of the promoter was clipped) for Cas9 expression. All have *nanos* 3′ UTR elements. At site B (275 bp from site A), the same *EGFP* gene is oriented in the opposite direction from Cas9. Eleven different Cas9 element variants are included, all of which use *nanos* regulatory elements. Some have an additional nuclear localization signal, while others have different *nanos* promoter elements, a Syn21 intron, additional 3′ UTR loop elements, an additional Kozak sequence, or a PEST sequence in Cas9.

A previous study examined the NosBN variant of *nanos*, that was truncated at the 5′ UTR by a large insertion, reported that it was still expressed in the germline, but not maternally deposited^44^. An initial experiment with an entirely truncated promoter using a complete drive with an EGFP target site showed no cleavage activity in females, while males retained 70% drive inheritance (compared to 80% with a complete *nanos* promoter) with very little germline resistance allele formation (Figure S2, Data Set S1). Following this observation, we truncated the *nanos* promoter to various lengths to determine if there is an optimal length with acceptable drive conversion in both sexes but no embryo resistance (Figure 1).

Additionally, we tested other variants of the Cas9 element. In a study of *Drosophila suzukii*, a combination of native (as opposed to *D. melanogaster*) *nanos* regulatory elements and the addition of a second nuclear localization signal (NLS) for Cas9 substantially increased drive conversion rate and total cut rate^23^. We thus created a construct with a second nuclear localization signal to examine this factor (though there were also silent changes to the coding sequence of our construct compared to our one nuclear localization signal standard version). Another reported method to increase expression is the use of a Syn21 intron^35^, which we also tested. A previous study examining the 3′ UTR of *nanos* found a stem-loop for interaction with Smaug and Oskar proteins, which competitively bind to *nanos* mRNA for degradation and translation, respectively, in nurse cells and the early embryo. We adjusted the 3′ UTR of *nanos* to provide three copies of this loop, seeking to increase mRNA degradation in the early embryo to reduce embryo resistance. As a method to increase Cas9 activity via increased translation, we inserted a *D. melanogaster* consensus Kozak sequence (AAAAATCAAA)^36^ between the standard 5′ UTR and Cas9 coding sequence. Finally, we used a variant with a PEST sequence, which we previously demonstrated was highly effective at reducing Cas9 activity due to reduced protein half-life^32^.

### Drive performance of truncated *nanos* promoter variants

Transgenic Cas9 flies with truncated *nanos* promoters were crossed to a split driving gRNA line targeting *yellow*. Females that were heterozygous for both the drive and Cas9 were then crossed to *w*^*1118*^ males, and progeny were phenotyped to assess drive performance. The drive inheritance rate was simply the fraction of flies inheriting the drive (gRNA) allele, marked by DsRed. Germline resistance allele formation could be detected by examining the fraction of male progeny that had the *yellow* knockout phenotype but not DsRed (though such alleles could have also formed if the allele remained uncut in the germline and was then cut by maternally deposited Cas9 and gRNA — however, this is less likely than formation in the germline due to generally higher germline cut rates). Embryo resistance allele formation was specifically measured in female offspring that also inherited the drive allele. For these, if maternally deposited Cas9 and gRNA cut the paternally inherited wild-type allele, a resistance allele would form, the female would show a yellow body color. These females could also be mosaic, indicating that maternally deposited Cas9 acted on only a fraction of the cells in the new offspring after the zygote had undergone one or more rounds of division. Note that only nonfunctional resistance alleles could be identified by yellow body color phenotype, but this represents a substantial majority of resistance alleles (perhaps over 90%^41,45,46^).

The full length (672 bp) *nanos* promoter yielded nearly 90% drive inheritance, but also 85% embryo resistance allele formation (Figure 2, Data Set S2). Variants were selected based on good PCR primer locations that cut the size successively by half, with a few additional variants selected near a critical region where performance rapidly changed. Variants with 305, 167, and 70 bp all had similar performance, indicating the most important enhancers for *nanos* are fairly close to the transcription start site. When the promoter was truncated to 59 bp, however, both drive conversion efficiency and embryo resistance allele formation were slightly decreased. At 48 and 37bp, embryo resistance was entirely abolished, and drive inheritance fell substantially, with many wild-type alleles remaining uncut in the germline. With no promoter, detectable Cas9 activity in females was abolished, consistent with our initial experiment.

**Figure 2.**
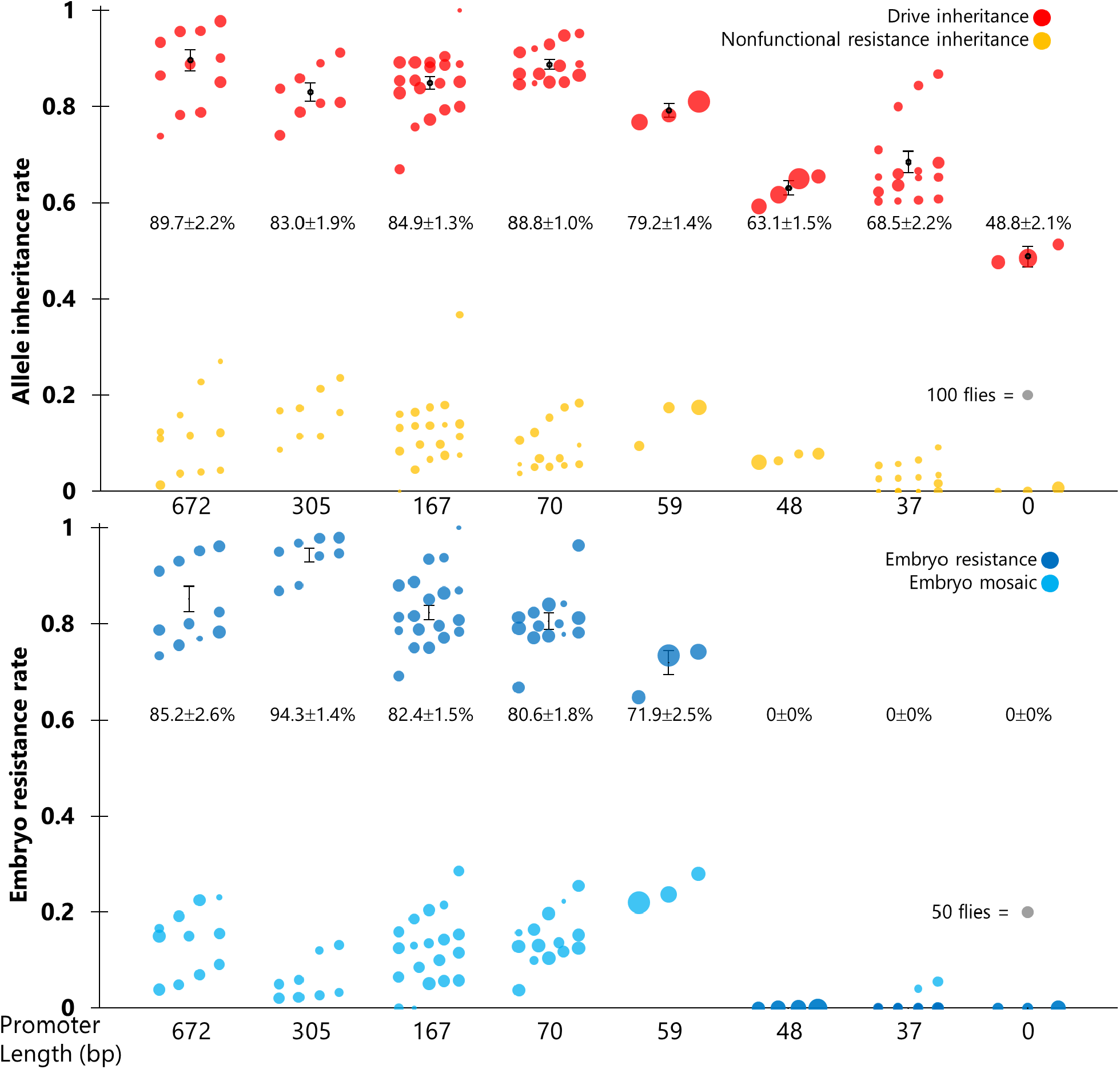
*nanos* promoter variants with different lengths modulate Cas9 homing drive activity. Drive inheritance of the split element targeting *yellow*. Cas9 elements had varying *nanos* promoter lengths before the 5′ UTR, with 672 bp being the standard length used in previous studies. Females that were heterozygous for both the drive and Cas9 (with a male Cas9 parent and female drive parent) were crossed to *w*^*1118*^ males, and their progeny were phenotyped. The drive inheritance rate is the percentage of these offspring with DsRed fluorescence. Germline nonfunctional resistance inheritance is only visible in male offspring (with yellow body but not DsRed phenotype). Embryo resistance rate is the proportion of DsRed female offspring with yellow body, and the mosaic rate is the fraction of yellow mosaic individuals within the DsRed females. Each dot represents the offspring from a single vial, and the size of the dots is proportional to the total number of offspring. Displayed values represent the average for drive allele inheritance and embryo resistance (± standard error of the mean).

Using JASPAR (https://jaspar.elixir.no/analysis), several transcription factors were identified that potentially bind to the 70 bp *nanos* promoter (CG4328, Ubx, BEAF-32, Dref, and Dbx), potentially explaining why it retains its activity. BEAF-32 and Dref belong to C2H2 zinc finger family and have a binding site motif (TATCGATA) between the 48 and 59 bp promoters. BEAF-32 mostly binds to the core regulatory region of highly active genes close to the transcription start site^47^, potentially accounting for the rapid loss of cleavage activity between the constructs with 48 and 59 bp promoters.

### Effect of other *nanos*-Cas9 element modifications on homing drive efficiency

To assess how other regulatory elements may manipulate drive performance, we constructed additional Cas9 elements with a variety of enhancements and used the same cross scheme with the split drive targeting *yellow*. First tested was a Cas9 construct with two nuclear localization signals (SV40 and nucleoplasmin), which was expected to have higher cleavage rates than our standard drive with only one nuclear localization signal (SV40). However, it only resulted in similar performance with perhaps slightly increased embryo resistance rate (Figure 3, Data Set S3). This result is not entirely unexpected because standard *nanos*-Cas9 already has very high cleavage rates both in the germline and embryo, leaving little room for increased cleavage.

**Figure 3.**
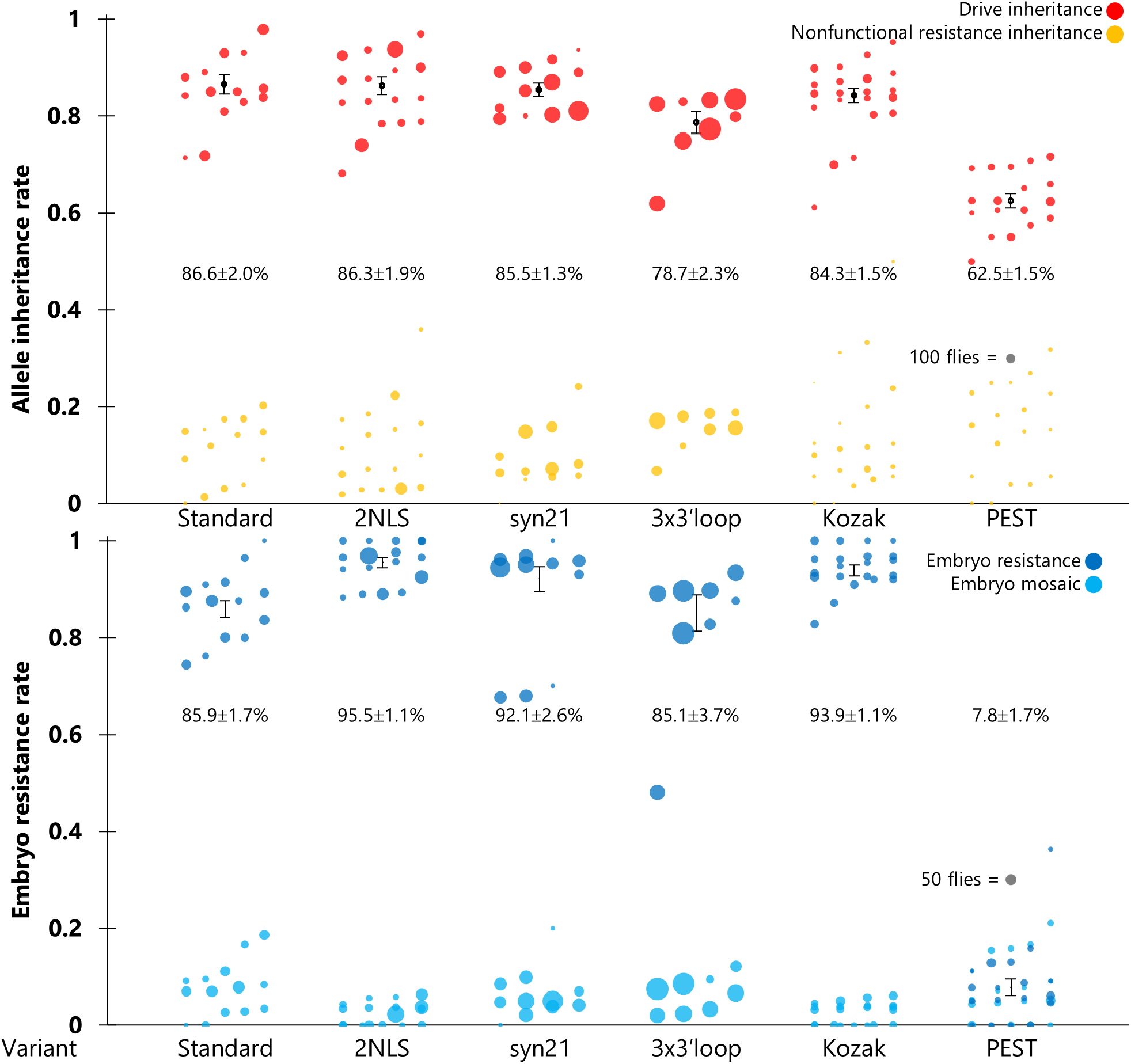
The effect on drive performance of different Cas9 gene modifications. Drive inheritance of the split element targeting *yellow*. Cas9 genes had two nuclear localization signals (NLS) in the Cas9 sequence, a Syn21 intron after the *nanos* 5′ UTR, additional repeats of a loop element (designed to repress translation in embryos) in the *nanos* 3′ UTR, an additional Kozak sequence before the coding sequences, or a PEST sequence at the C-terminus of the Cas9 gene. Females that were heterozygous for both the drive and Cas9 (with a male Cas9 parent and female drive parent) were crossed to *w*^*1118*^ males, and their progeny were phenotyped. The drive inheritance rate is the percentage of these offspring with DsRed fluorescence. Germline nonfunctional resistance inheritance is only visible in male offspring (with yellow body but not DsRed phenotype). Embryo resistance rate is the proportion of DsRed female offspring with yellow body, and the mosaic rate is the fraction of yellow mosaic individuals of all DsRed females. Each dot represents the offspring from a single vial, and the size of the dots is proportional to the total number of offspring. Displayed values represent the average for drive allele inheritance and embryo resistance (± standard error of the mean).

For the Syn21 intron construct and consensus Kozak sequence construct, performance was even more similar to the standard *nanos*-Cas9 (Figure 3, Data Set S3). However, for the 3x3′loop construct that contained three repeats of a 180 bp segment of the *nanos* 3′ UTR, drive inheritance was somewhat lower, while embryo resistance remained the same as our standard construct. Although this construct was designed to suppress translation in the early embryo, it is possible that embryo resistance occurs almost immediately as a result of deposited Cas9 protein, making new translation less important. The reduction in drive inheritance could potentially be explained by the large repeated element. A different type of repeated element was shown to drastically affect homing drive inheritance in another study^48^. The final variant tested contained a PEST sequence that acts as a signal for protein degradation and thus should reduce Cas9 half-life, which a previous study showed would usually abolish most drive activity in constructs with intermediate activity, but only marginally affect a high activity system. With the highly active *nanos* promoter, some activity remained, but drive inheritance was reduced without a corresponding reduction in the germline resistance rate. This suggests that at least a moderate proportion of resistance alleles may form early, before the PEST sequence takes effect. Embryo resistance was also nearly eliminated with this construct.

### Combinations of regulatory elements modulate drive inheritance and resistance allele formation

We next sought to combine regulatory elements to further optimize drive activity and assess which combinations may be compatible. First, we used repeats of promoter elements, both of the 70 bp and 48 bp promoters. With three repeats of the 70 bp promoter, drive inheritance was similar to our test with one 70 bp promoter element (see Figure 2), but embryo resistance fell markedly in about half of individuals tested, while remaining high in the other half (Figure 4, Data Set S4). It is unclear why embryo resistance was only lower in a fraction of flies, despite having more promoter elements. It is possible that adjacent enhancers may have created transcriptional interference, reducing transcription just enough to affect embryo resistance but not germline cleavage. When we tested the construct with three repeats of the 48 bp promoter, cleavage activity was substantially increased compared to the system with a single 48 bp promoter and was similar to the complete promoter, but embryo resistance was also raised back to the high level of the standard *nanos* promoter. This indicates that smaller repeated regions are unlikely to significantly impair drive performance and that repeating a promoter element can substantially increase cleavage rates in the context of gene drives.

**Figure 4.**
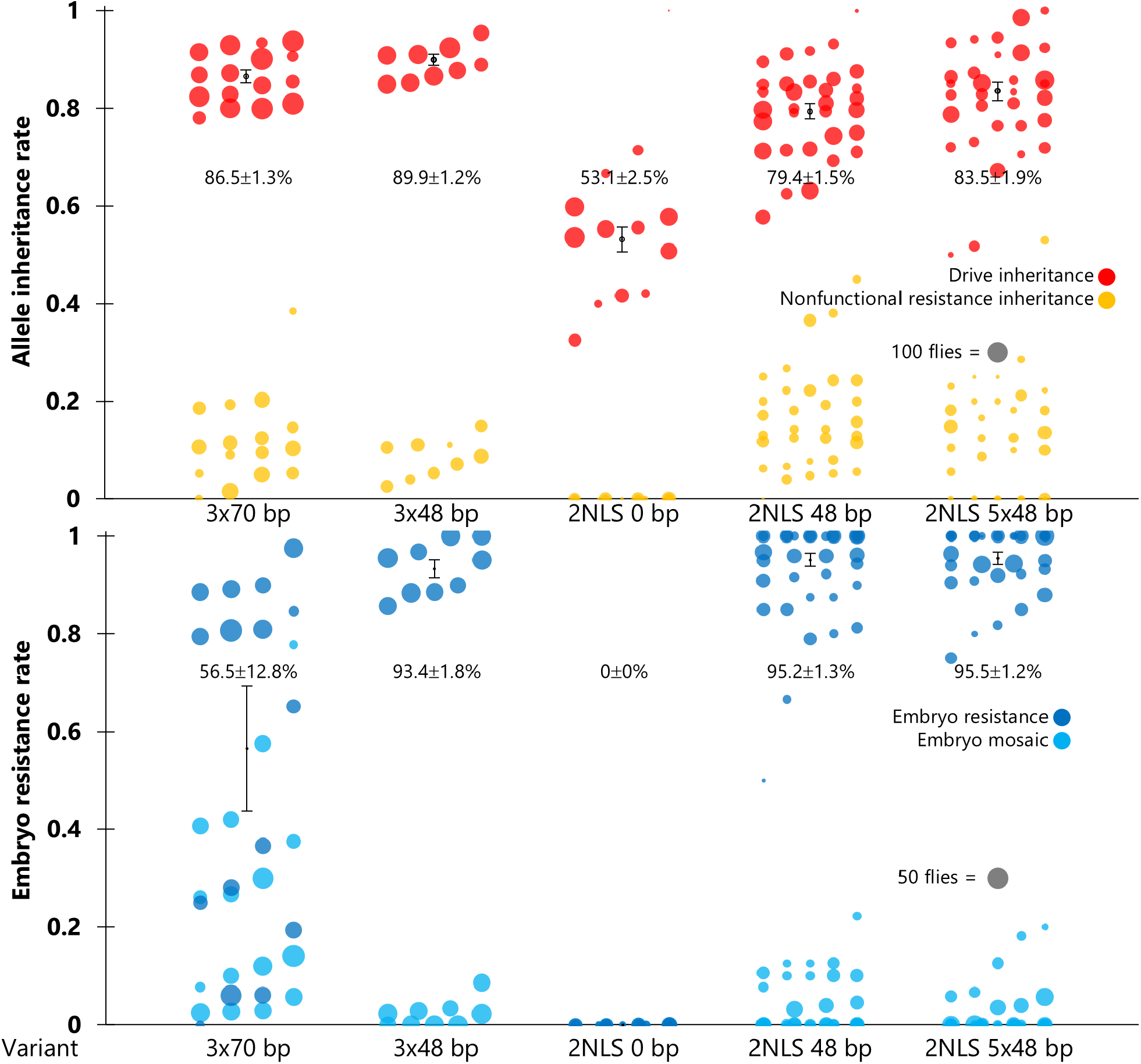
Effects of Cas9 element variant combinations. Drive inheritance of the split element targeting *yellow*. Some Cas9 genes have three repeats of a 70 bp promoter, or three repeats of a 48 bp promoter. Others have two nuclear localization signals (NLS) combined with no promoter before the *nanos* 5′ UTR, a 48 bp promoter element, or five repeats of a 48 bp promoter element. Females that were heterozygous for both the drive and Cas9 (with a male Cas9 parent and female drive parent) were crossed to *w*^*1118*^ males, and their progeny were phenotyped. The drive inheritance rate is the percentage of these offspring with DsRed fluorescence. Germline nonfunctional resistance inheritance is only visible in male offspring (with yellow body but not DsRed phenotype). Embryo resistance rate is the proportion of DsRed female offspring with yellow body, and the mosaic rate is the fraction of mosaic individuals of the DsRed females. Each dot represents the offspring from a single vial, and the size of the dots is proportional to the total number of offspring. Displayed values represent the average for drive allele inheritance and embryo resistance (± standard error of the mean).

We also tested the use of two nuclear localization signals combined with 0 and 48 bp promoters. Ideally, this strategy would produce a germline-specific expression pattern that raises the germline cleavage rate, while keeping the embryo resistance rate at zero. However, no activity was measured with the 0 bp promoter, and performance with the 48 bp promoter again reverted to approximately the level of the standard *nanos* promoter (Figure 4, Data Set S4). This indicates that truncated *nanos* promoters probably only change the quantity of expression and may have a more limited effect on the timing of expression. A final test with two nuclear localization signals and five repeats of the 48 bp promoter yielded similar performance.

### Somatic cell cleavage appears in high activity *nanos*-Cas9 elements

While higher germline activity may be desirable in a wide variety of scenarios, potentially correlated somatic expression could harm efforts to build suppression drives with low fitness costs. One positive aspect of the standard *nanos* promoter is that somatic expression is minimal, but this may not be the case in constructs with higher expression. To assess this, we generated females that were heterozygous for both the gRNA element targeting *yellow* and Cas9 (inheriting Cas9 from the male parent) and determined the proportion of offspring that displayed mosaic body phenotype. We found that the promoter version with two nuclear location signals showed a small amount of mosaicism, indicating detectable somatic expression (Data Set S5). Because Cas9 was most likely expressed in the same spatial and temporal pattern, only varying in expression amplitude, it indicates that *nanos* does likely have some leaky somatic expression, just normally not high enough to induce a phenotype. Further, because mosaicism was seen in under 10% of females and represented no more than 20% of the body, it is unclear if this would be sufficient to induce a considerable fitness cost in suppression drives. Parallel testing of this construct with a drive targeting *doublesex* indicated no substantial visual phenotypic change (which was seen when somatic expression was higher^49^) compared to our standard *nanos*-Cas9 construct^50^. Our *nanos* variant with three copies of the 48 bp promoter produced no signs of somatic activity, nor did our version with two nuclear localization signals and a single 48 bp promoter. However, our variant with two nuclear localization signals and five copies of the 48 bp promoter did produce somewhat more mosaic females than the standard *nanos* promoter with two nuclear localization signals (Data Set S5).

### Performance comparisons using other gRNA elements

While it is difficult to distinguish higher Cas9 activity compared to standard *nanos*-Cas9 in the drive targeting *yellow*, this may be possible in other systems with naturally lower cleavage rates. One of these involves a split homing rescue drive targeting the haplolethal gene, *RpL35A*. In this system, two gRNAs are linked by tRNAs^22^. The drive has a functioning copy of the target gene, but any genotype with a nonfunctional resistance allele is nonviable (Figure S1). When together with standard *nanos*-Cas9, drive inheritance in males and females is approximately 90%, and females have approximately 30% fewer offspring, mostly due to embryo resistance^22^.

When tested with the two nuclear localization signal variant, drive inheritance in the offspring of male heterozygotes was somewhat increased to 96% (Figure 5, Data Set S6). However, drive inheritance from female carriers substantially declined. This may be related to an even more substantial decline in offspring viability. In our protocol, parents were allowed to lay eggs for a week, which was usually enough to saturate the vial and provide many offspring. However, in all cases with the *RpL35A* drive in females, only a tiny fraction of progeny were recovered, with some dead larvae and pupae observed. This was likely due to much higher embryo resistance (and incomplete mosaic resistance), forming many lethal resistance alleles in the progeny of females. In cells experiencing drive conversion, there may have been a higher amount of Cas9 and thus higher mortality in the offspring derived from these cells, thus enriching for offspring of cells with abnormally low Cas9 expression (accounting for lower drive inheritance). It is also possible that tissues with lower Cas9 expression may have had increased health and given rise to more offspring that were the product of less drive conversion.

**Figure 5.**
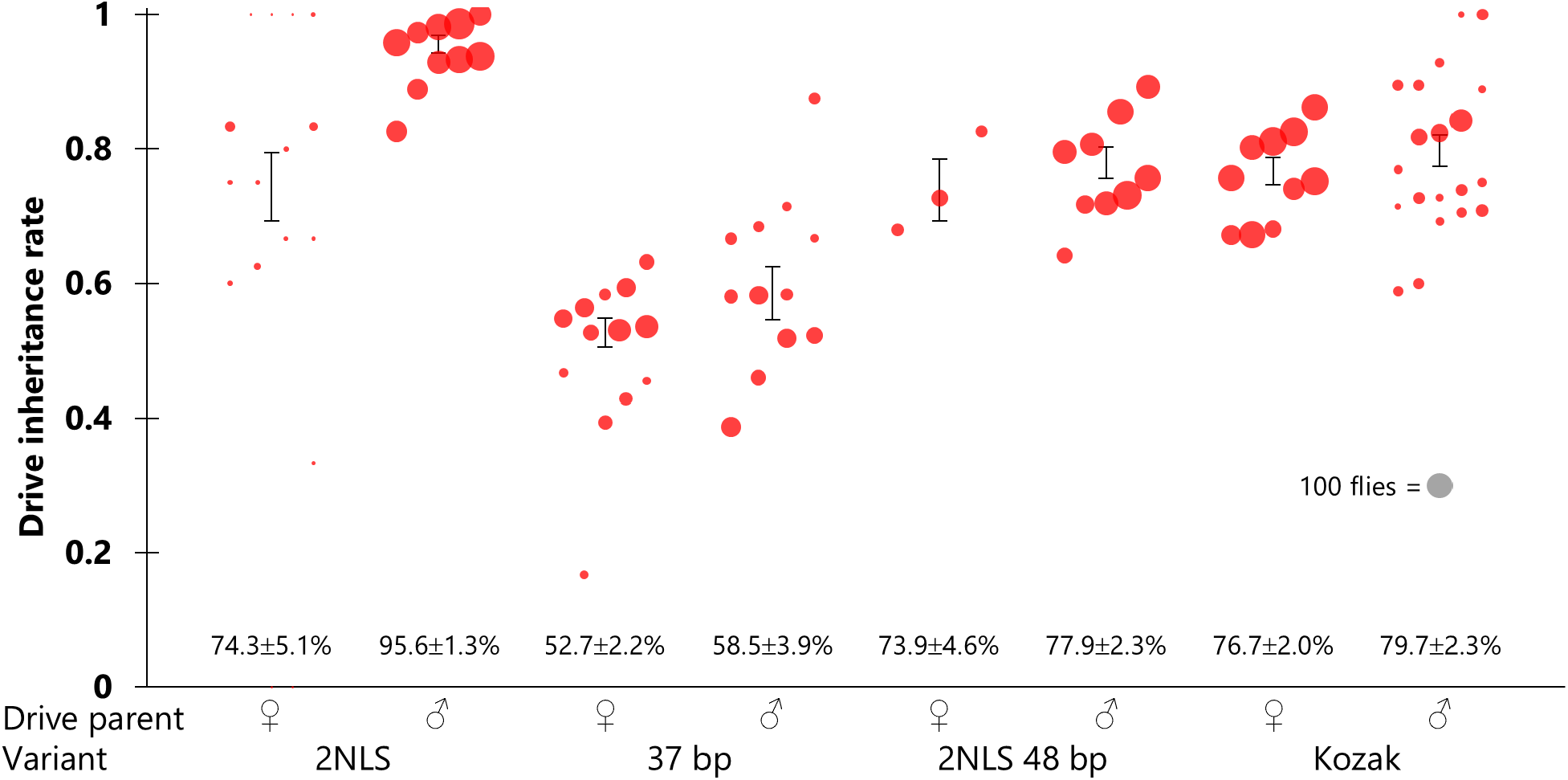
Cas9 element variant performance with the homing rescue drive. Drive inheritance of the split element that targets *RpL35A* (a haplolethal gene) while providing rescue. Cas9 genes had two nuclear localization signals, a 37 bp *nanos* promoter element, two nuclear localization signals (NLS) combined with a 48 bp promoter element, or a *Drosophila* consensus Kozak sequence. Flies that were heterozygous for both the drive and Cas9 (with a male Cas9 parent and female drive parent) were crossed to *w*^*1118*^, and their progeny were phenotyped. The drive inheritance rate is the percentage of these offspring with DsRed fluorescence. Individuals with a nonfunctional resistance allele (and many that are mosaic for such alleles) were nonviable. Each dot represents the offspring from a single vial, and the size of the dots is proportional to the total number of offspring. Displayed values represent the average for drive allele inheritance (± standard error of the mean).

In another test, we assessed the 37 bp Cas9 element, finding low drive inheritance among males and even lower among females (Figure 5, Data Set S6), consistent with another study where several constructs had higher Cas9 cleavage in males^51^. These results are also consistent with the notion that the *yellow* drive has intrinsically higher activity than the drive targeting *RpL35A*, possible due to gRNA sequence activity, the genomic location affecting gRNA expression, or the use of tRNAs in the *RpL35A* system. We also tested the 48 bp promoter with two nuclear localization signals and the consensus Kozak sequence that should increase Cas9 translation.

Both yielded intermediate performance, worse than the standard *nanos*-Cas9. For the former, the reduced activity of the *RpL35A* drive may have allowed the reduced activity of the Cas9 element to be more easily measured. However, for the consensus Kozak element, it is not clear why it lost activity compared to standard *nanos*-Cas9. It is possible that the Kozak element interacts with other sequence elements to start translation, so despite being a theoretically optimal element for *Drosophila melanogaster*, gave inferior performance to the natural Kozak element of *nanos* (this *nanos* sequence was still present, but may not have functions due to interruption by the added sequence or greater distance from the start codon).

Similarly, we conducted some tests with a split homing suppression element targeting the haplosufficient female fertility gene *yellow-G* with four gRNAs (linked by tRNAs) ^43^. This one is thought to have intermediate activity, in between the homing rescue drive and the drive targeting *yellow*^32^. Unexpectedly, the Cas9 with two nuclear localization signals decreased both male and female drive inheritance by about 10% (Figure S3, Data Set S7) compared to a previous study with the standard *nanos*-Cas9 element^43^. The reasons for this result are unclear. For females, it may have been related to loss of fertility in some tissues with high drive activity, leading to overrepresentation of offspring from a minority of remaining tissues with lower cut rates (somewhat similar to what may have caused reduced performance in the homing rescue drive). For males, it is possible that higher Cas9 activity in early stages of the germline led to more resistance alleles forming before there was a chance for drive cleavage, though this is contrary to the pattern observed in the *yellow* drive. We also tested the 37 bp promoter, which showed some activity in males (less than the drive targeting *yellow*, but slightly more than the drive targeting *RpL35A*), but none in females (Figure S3, Data Set S7).

## Discussion

In this study, we explored the effect of modifications to the *nanos* promoter, 5′ UTR, 3′ UTR, and Cas9 itself. Though the *nanos* promoter has low somatic expression, making it a desirable starting point for gene drive optimization, we found no combination that could both preserve high germline cut rates yet also substantially reduce maternal Cas9 deposition. Because genes like *nanos* and *vasa* are highly expressed in the germline and required in the embryo, perhaps these promoters are inextricably linked to both processes. However, we did successfully reduce and increase Cas9 activity, showcasing several potentially useful options for improving gene drives in a wide range of species. Indeed, previous studies that tested gene drives with meiosis-specific promoters *Stra8* and *Spo11* in mice found that expression levels were either altogether too low (*Spo11*)^26^, or likely too high and too early in the germline (*Stra8*)^52^. These promoters could potentially benefit from modifications as demonstrated here.

A recent study examining many different promoters for Cas9 in the context of a homing gene drive found that near-optimal expression is possible (with high drive conversion and low embryo resistance), but that this likely involved an extremely narrow window for the optimal quantity of expression in the female germline that may be very difficult to achieve^51^. Nevertheless, by gaining additional tools to increase or decrease expression, there may be more opportunities in the future to achieve optimal expression, particularly in non-model organisms where high performance is required for release candidates (especially in suppression drives).

Unlike the other adjustments we considered, the 2xNLS is likely the simplest in terms of its effect. It only slightly changes the protein sequence of Cas9 and is thus unlikely to affect the specific expression pattern or level of expression. However, it should result in a greater fraction of Cas9 being localized to the nucleus and thus increase Cas9 activity. This resulted in several interesting observations. First, when combined with the standard *nanos* promoter, some somatic activity was observed, despite *nanos* being a well-known germline-specific promoter (substantially more germline specific than others in a recent promoter comparison study^32^). This may imply that almost any gene with sufficiently high expression in the germline may have some residual expression in somatic cells. If this is the case, it would mean that a potential universal solution for somatic expression may be to reduce overall activity levels via any mechanism possible. In this case, the ideal outcome would be a reduction of somatic expression but retention of high germline cleavage activity (and therefore high drive efficiency).

Another interesting observation with the 2xNLS line was that drive efficiency often changed, even when cleavage rates were already near-100% with a single NLS. This may be due to complexities of the target site and indicate that in some situations, higher activity may not be desirable, even if embryo resistance or somatic activity could be tolerated.

Related to this is the poor performance of the haplolethal-targeting homing rescue drive with the 2xNLS Cas9. When combined with the standard Cas9, the level of embryo resistance (based on nonviable offspring) is about 30%. However, nearly 100% of offspring were nonviable with the 2xNLS version. This implies a greater than 3-fold increase in the level of embryo resistance, which means a greater than 3-fold increase in the Cas9 activity level. However, it is possible that the increase in activity level was lower, because the exact conditions in which offspring are nonviable is not clear. While a full embryo resistance allele would usually fill this criterion (with perhaps some exceptions in certain genetic backgrounds^53^), it is likely that a sufficiently high level of mosaicism would also render most individuals nonviable. Passing this threshold may not require a greater than 3-fold increase in activity. On the other hand, the level of embryo resistance with the truncated *nanos* promoter went from 0% (without any mosaicism) to 90% when targeting the *yellow* gene, which implies an increase in activity level far higher than 3-fold. Similarly, the appearance of somatic activity when none existed before is also potentially indicative of a greater increase in activity. These results contrast with non-drive studies showing more modest improvements by the addition of a second NLS. The cause of this discrepancy remains unclear, though it may be related to genomic expression in our system and cell-based experiments in many other systems. Perhaps other factors are involved in genomic transcription, translation, or other processing of Cas9 with an additional NLS.

Using a *nanos*-Cas9 element as a basis for investigation, we demonstrated options for increasing or reducing expression levels of Cas9-based gene drives using promoter truncations, copying of promoter elements, and nuclear localization signals. By applying combinations of these techniques, it may be possible to gain finer control of Cas9 activity to achieve improved gene drive performance. Future investigations in non-model organisms in particular my benefit from these types of optimizations after initial assessments of drive performance.

## Supporting information

Supplemental Data Sets

## Acknowledgements

This study was supported by the Center for Life Sciences and the National Natural Science Foundation of China (grant 32270672). JD was supported in part by the Postdoctoral Fellowship of Peking-Tsinghua Center for Life Sciences. NRF gratefully acknowledges the Dutch graduate school for Production Ecology & Resource Conservation for funding.

## Supplemental Information

**Figure S1.**
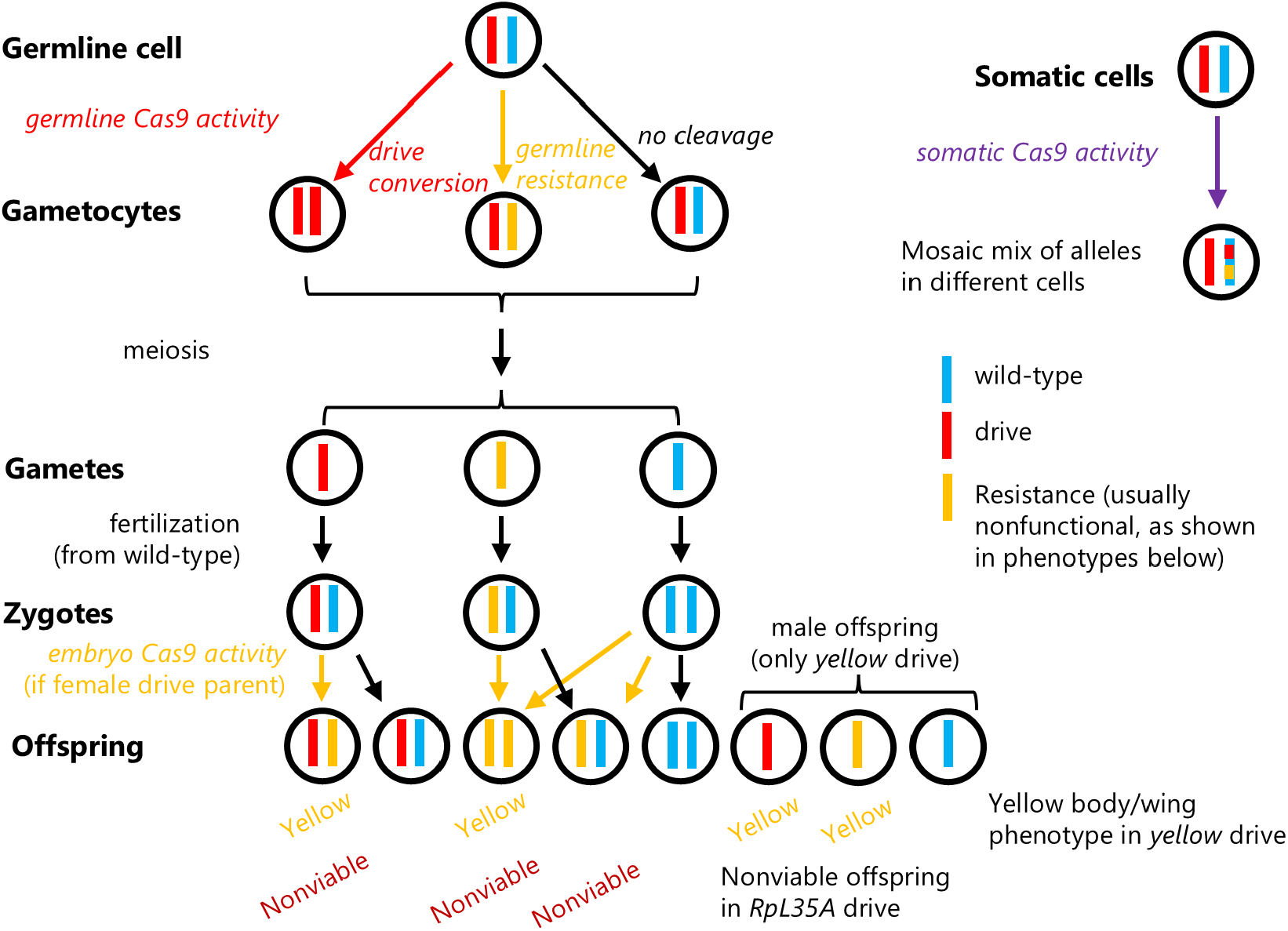
Genotypes resulting from drive activity. Drive conversion occurs in germline cells of drive/wild-type heterozygotes via homology-directed repair. However, resistance allele formation by end-joining can also occur at this stage, and some wild-type alleles may remain uncut. In new zygotes and early embryos, cleavage from maternally deposited Cas9 and gRNA can form resistance alleles (this can be mosaic if it occurs shortly after the zygote stage, leading to different genotypes in different cells). The drive targeting *yellow* is X-linked, and males will thus only have a single allele. Any individual lacking a wild-type allele or a functional resistance allele will have a yellow body color. For the drive targeting the haplolethal gene *RpL35A*, drive and wild-type alleles both have a functioning copy of the gene, but any individual with even one nonfunctional resistance allele will be nonviable. Rare functional resistance alleles have the same phenotype as wild-type alleles in these drives. Somatic Cas9/gRNA expression can change the genotype in some cells of drive/wild-type heterozygotes, resulting in mosaic genotypes that are visible in individuals with the *yellow* drive.

**Figure S2.**
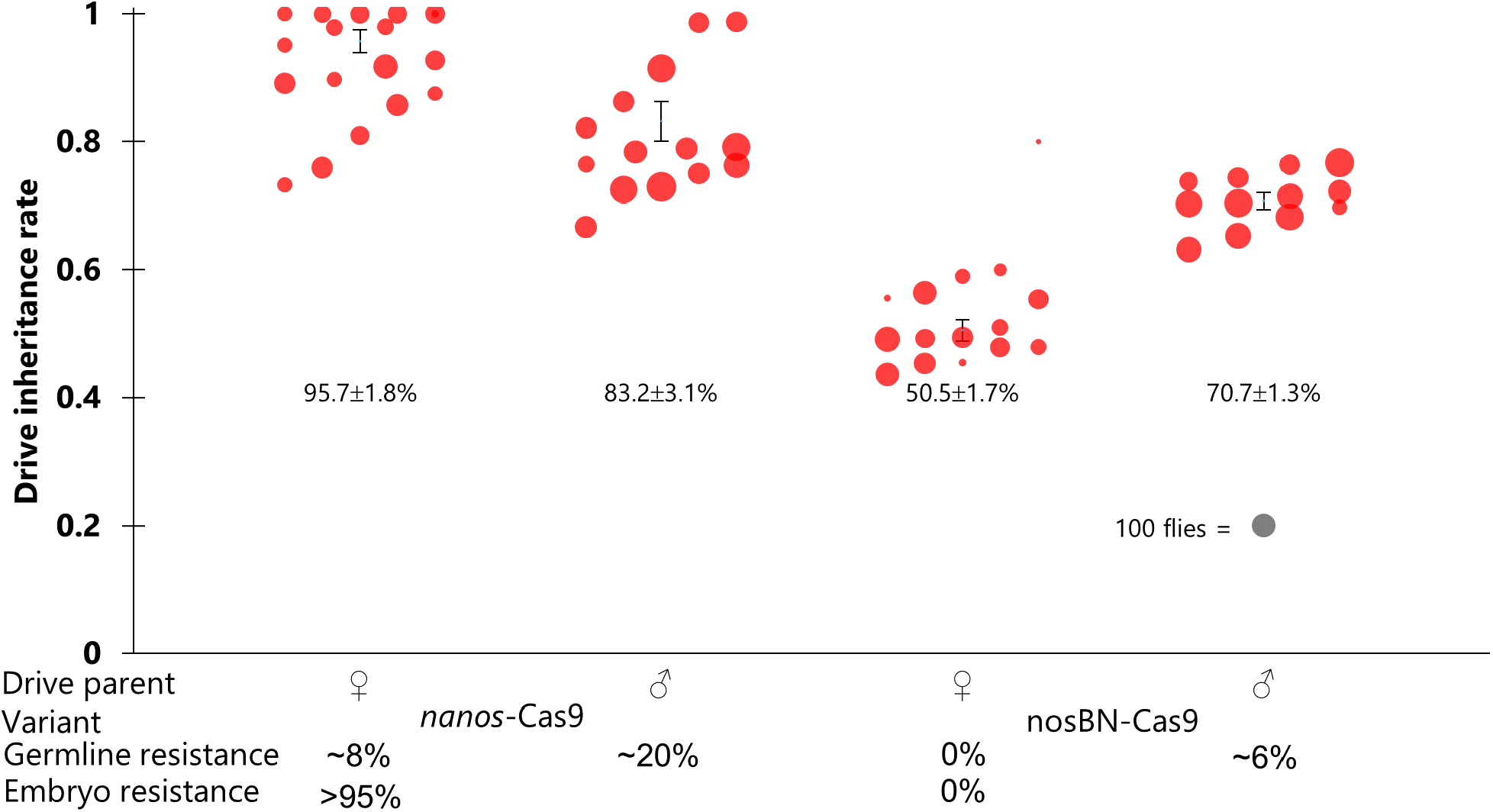
Effect of *nanos* promoter removal on a drive targeting EGFP. Drive inheritance of complete drives targeting EGFP. Flies that were heterozygous for both the drive and Cas9 (with a male drive parent and female EGFP parent) were crossed to EGFP homozygotes or *w*^*1118*^, and their progeny were phenotyped. The drive inheritance rate is the percentage of these offspring with DsRed fluorescence. Each dot represents the offspring from a single vial, and the size of the dots is proportional to the total number of offspring. Displayed values represent the average for drive allele inheritance (± standard error of the mean). nosBN is a variant that lacks the *nanos* promoter but retains the nanos 5′ UTR (and some sequence from an adjacent P element). Data for the first drive is from a previous study.

**Figure S3.**
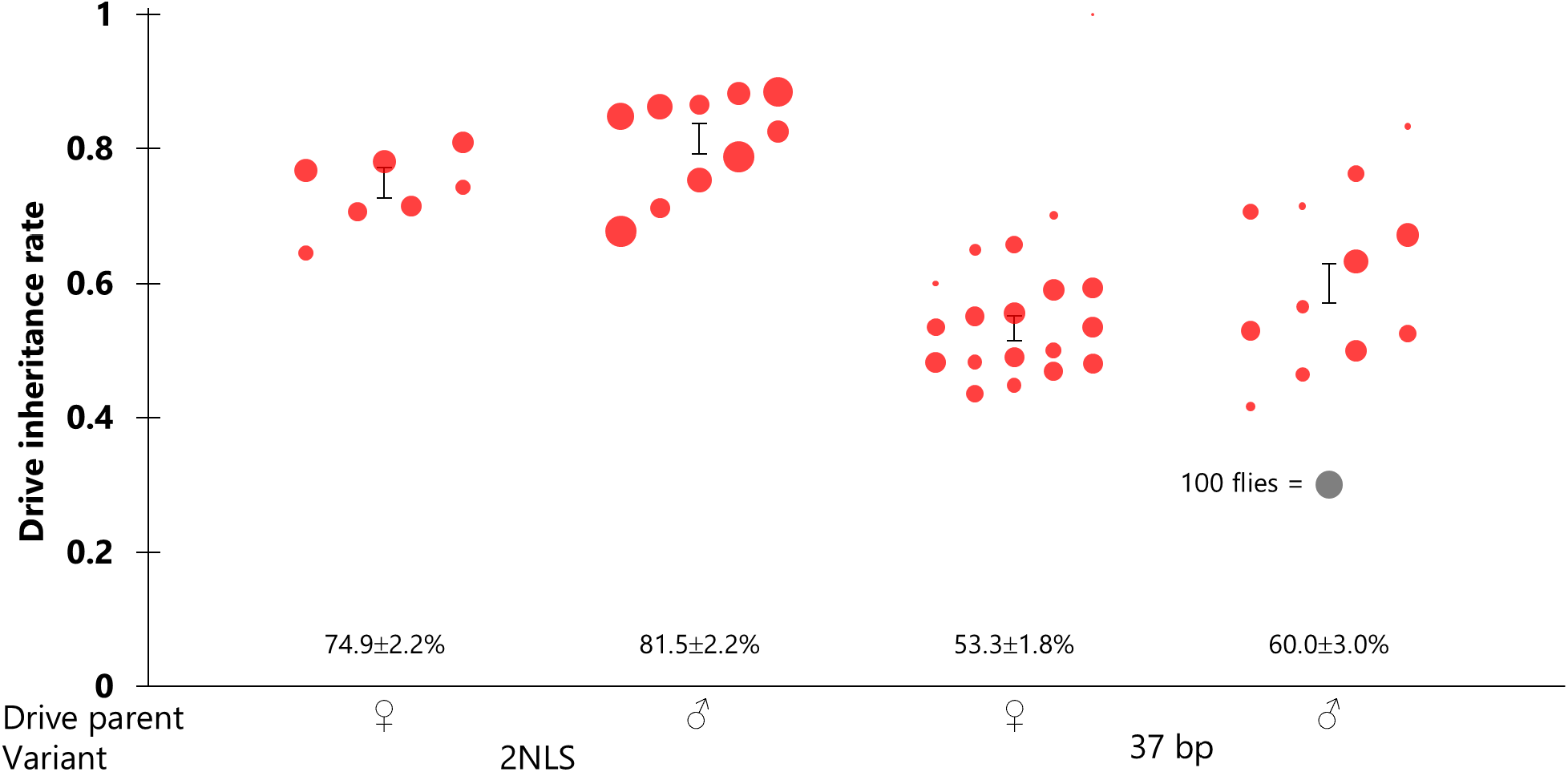
Drive performance with the suppression drive. Drive inheritance of a split suppression element targeting *yellow-G* (a haplosufficient female fertility gene). Flies that were heterozygous for both the drive and Cas9 (with a male Cas9 parent and female drive parent) were crossed to *w*^*1118*^, and their progeny were phenotyped. The drive inheritance rate is the percentage of these offspring with DsRed fluorescence. Each dot represents the offspring from a single vial, and the size of the dots is proportional to the total number of offspring. Displayed values represent the average for drive allele inheritance (± standard error of the mean).

